# ContaTester: Fast cross-contamination estimation and identification for large human sequencing cohorts

**DOI:** 10.1101/2021.10.01.461647

**Authors:** Damien Delafoy, Jonathan Mercier, Elise Larsonneur, Nicolas Wiart, Florian Sandron, Thomas Mejean, Stéphane Meslage, Delphine Daian, Robert Olaso, Anne Boland, Jean-François Deleuze, Vincent Meyer

**Affiliations:** Université Paris-Saclay, CEA, Centre National de Recherche en Génomique Humaine, Evry, France

**Keywords:** contamination, VCF, allelic balance, whole genome sequencing

## Abstract

**Background:** Interest in genomic medicine for human health studies and clinical applications is rapidly increasing. Clinical applications require contamination-free samples to avoid misleading results and provide a sound basis for diagnosis.

**Results:** Here we present ContaTester, a tool which requires only allele balance information gathered from a VCF file to detect cross-contamination in germline human DNA samples. Based on a regression model of allele balance distribution, ContaTester allows fast checking of contamination levels for single samples or large cohorts (less than two minutes per sample). We demonstrate the efficiency of ContaTester using experimental validations: ContaTester shows similar results to methods requiring alignment data but with a significantly reduced storage footprint and less computation time. Additionally, for contamination levels above 5%, ContaTester can identify contaminants across a cohort, providing important clues for troubleshooting and quality assessment.

**Conclusions:** ContaTester estimates contamination levels from VCF files generated from whole genome sequencing normal sample and provides reliable contaminant identification for cohorts or experimental batches.

## Introduction

Advances in genomic medicine are steadily improving and facilitating diagnosis and healthcare. In parallel, a need is arising to develop robust quality control methods to ensure reliable contamination-free data. Sources of contamination can differ, ranging from human and technological errors to bioinformatics processing errors, consequently the contamination risk needs to be carefully assessed. To evaluate contamination levels, current tools use two main sources of evidence: Sequence Alignement Map (SAM/BAM) and Variant Call Format (VCF).

Most tools developed for contamination detection are dedicated to the somatic context and require the use of paired samples to determine contamination levels; these tools include Conpair [1], GATK CalculateContamination [2] and HYSYS [3]. To our knowledge, there are few tools dedicated to germline single sample contamination analysis. Common practice in the community is to use VerifyBamID2 [4], which requires BAM files as input to analyze mixture models, thus estimating the likelihood of contamination. ART-Deco [5] uses coverage tables provided by GATK [6] and analyzes the fraction of supporting reads to detect contamination in the restricted context of high-coverage panel analysis. Other programs like Peddy [7] use pedigree information to identify swaps or contamination within families.

Here we present ContaTester [8], an integrated solution for fast contamination estimation and contaminant identification in a Whole Genome Sequencing (WGS) normal sample. ContaTester is based on Allele Balance (AB) regression models and correlations, and requires only VCF including the Allele Depth (AD) metric; no other information is needed. In addition, we have developed a function for contaminant identification to support troubleshooting and quality control performed on a production platform.

## Methods

### Experimental and *in silico* design

Five DNA sample mixtures (1%, 5%, 10%, 15%) from both NA10859 and NA12878 individuals were experimentally designed and sequenced to a 30x depth of coverage to validate our method [9]. All samples were processed with a “Truseq™ PCRfree kit” and sequenced with a HiSeqX Illumina^®^ sequencer.

We selected sequencing datasets from two distinct families (CEPH/UTAH PEDIGREE 1463 and 1347) to simulate *in silico* mixtures from related and unrelated samples mimicking contamination conditions commonly observed in large cohorts. The mixtures included selected read ratios from NA12878, NA12891 (NA12878 father), NA12892 (NA12878 mother) and NA10859 (unrelated sample) (Table 1).

**Table 1.** Dataset is composed of 188 cases of contamination: 23 levels (0.5; 1.0; 1.5; 2.0; 2.5; 3.0; 3.5; 4.0; 4.5; 5.0; 7.5; 10.0; 12.5; 15.0; 17.5; 20.0; 22.5; 25.0; 30.0; 35.0; 40.0; 45.0; 50.0) x 8 combinations and 4 contamination’s free

|  | NA10859 | NA12878 | NA12891 | NA12892 |
| --- | --- | --- | --- | --- |
| NA10859 | - | 23 levels | 23 levels | 23 levels |
| NA12878 | 23 levels | - | - | - |
| NA12891 | 23 levels | - | - | 23 levels |
| NA12892 | 23 levels | - | 23 levels | - |

### Allele balance ranges and regression models

To determine the AB (1), we used the ratio of the number of reads supporting the first alternative allele, divided by the sum of numbers of reads supporting the reference allele with the first two alleles. Insertions-Deletions (InDels) were discarded from the Allele Balance distribution.

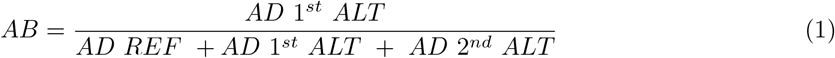

We evaluated several scenarios of AB range selections for contamination characterization. The mixture of two samples with different genetic backgrounds produced an increase in the number of variants, a spread of the allele balance distribution for ranges between 0.25 and 0.75 and peaks in low [0-0.25] and high [0.75-1] AB ratios (Figure 1).

**Figure 1.**
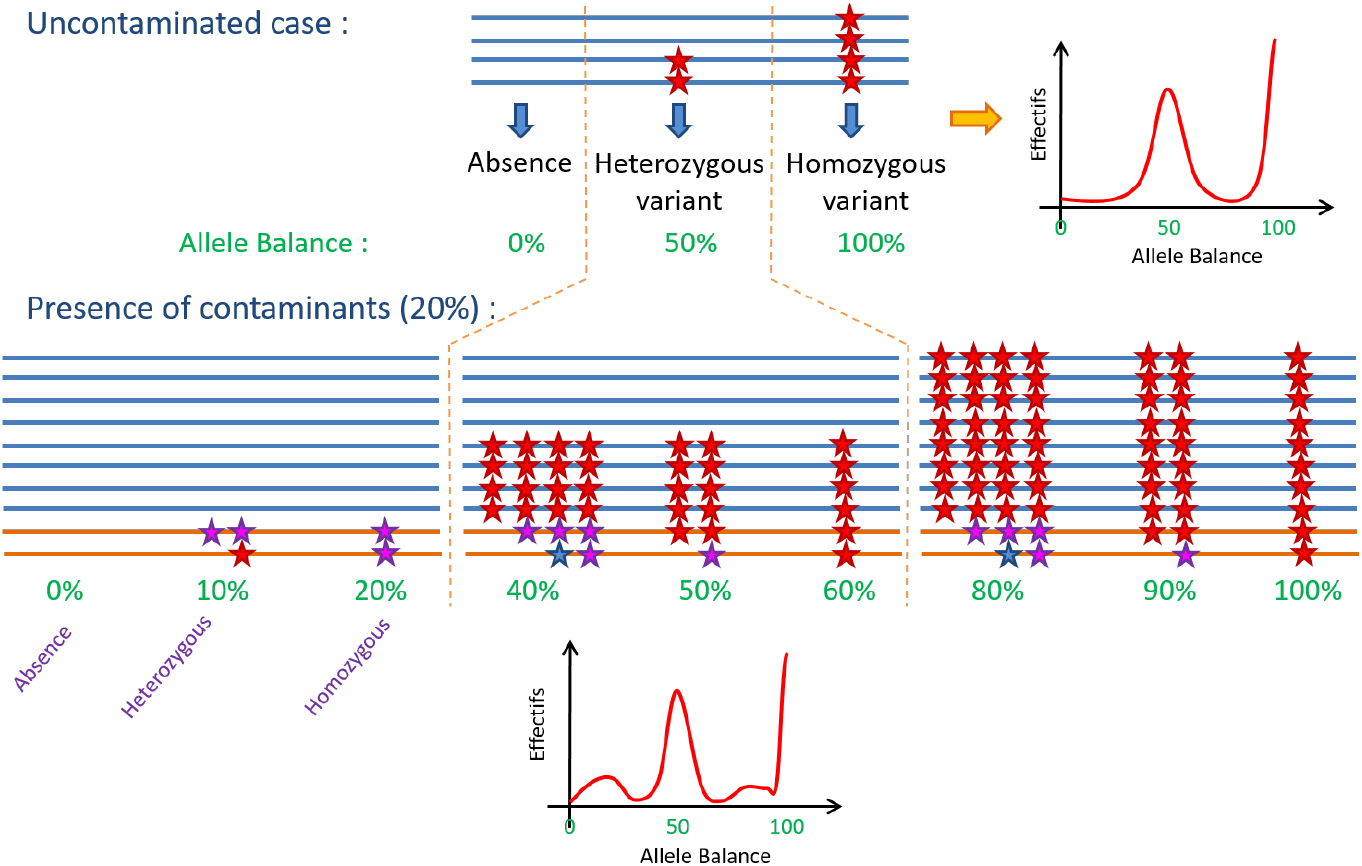
Schema of a theoretical 20% contamination impact on allele balance distribution. Blue lines represent reads alignments from initial sample. Orange lines represent reads alignments from contaminant. Stars represent different scenarios of single nucleotide variation

We selected the AB ranges between [0.18-0.49] and [0.51-0.82] gathering the highest numbers of variant according to the drift of their AB ratio for a second order polynomial regression (Figure 2). These two intervals maximize the coefficient of determination (*r*^2^)(Figure 3). The polynomial regression models had to be customized to fit depth of coverage conditions. Three regression models for common depth of coverage in WGS (30x, 60x and 90x) were calculated. Each condition showed a high *r*^2^ (*r*^2^ > 0.999) with the computed polynomial models.

**Figure 2.**
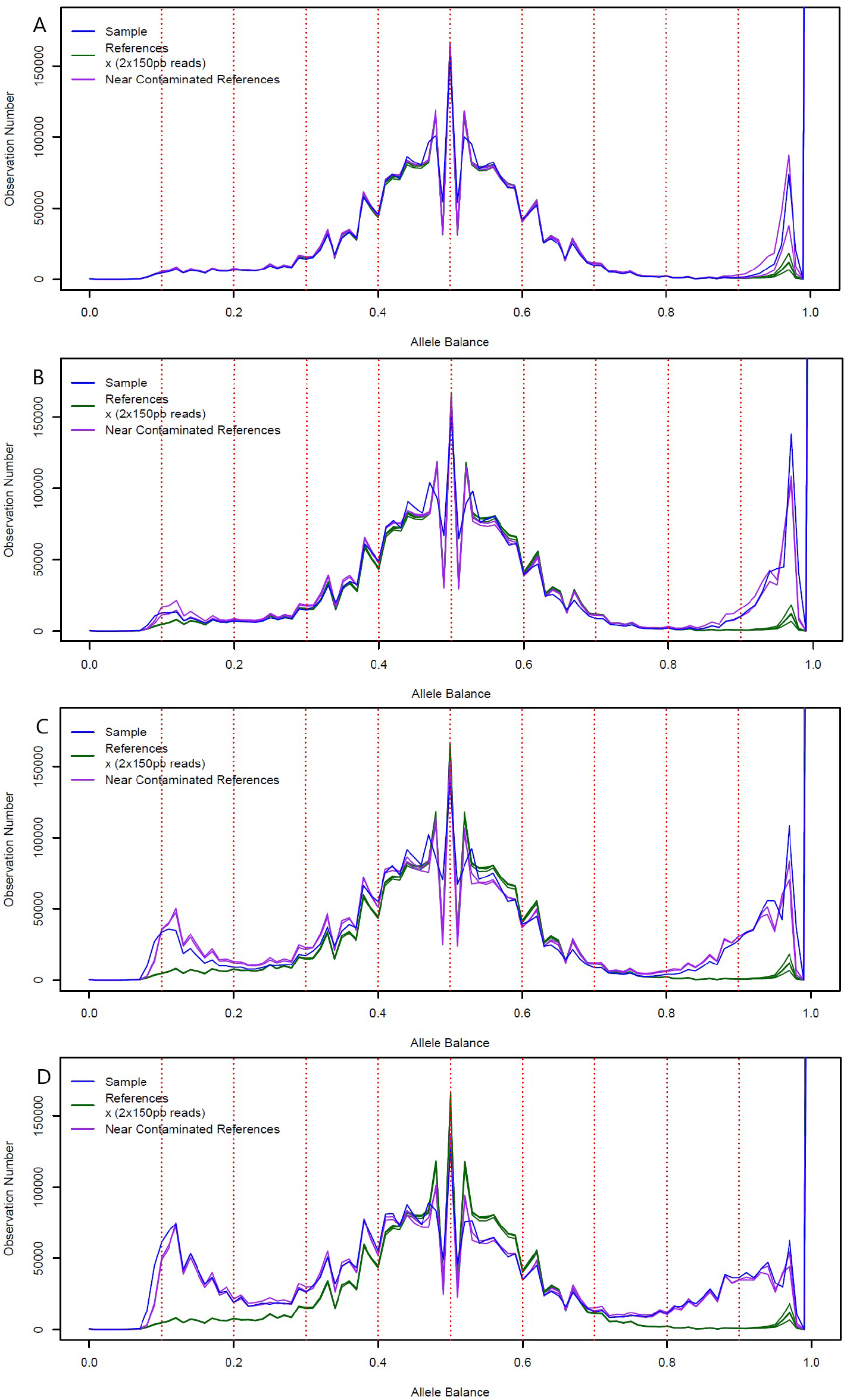
Allele balance distribution of experimentally contaminated sample (blue) versus four controls (green) and two in-silico contaminated sample (purple) for several ratio of mixture: A) NA12878 contaminated with 1% of NA10859 B) NA12878 contaminated with 5% of NA10859 C) NA12878 contaminated with 10% of NA10859 D) NA12878 contaminated with 15 % of NA10859

**Figure 3.**
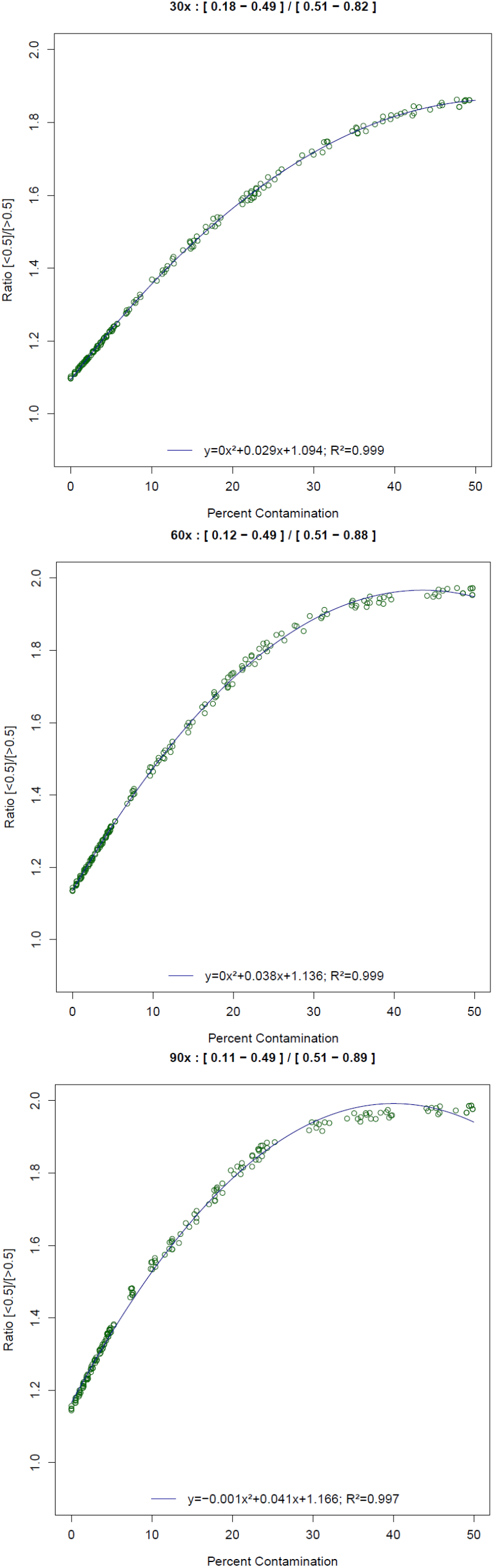
Polynomial regression of variant count ratio in selected allele balance range for contamination estimation at 30x, 60x and 90x

Additionally, we selected the ranges [0.01-0.49] and [0.51-0.99], removing the expected common AB peaks to perform a Pearson’s correlation analysis as a complementary method to support the contamination identification.

### Contaminant research and identification

For this study, we selected variants associated with an AB range between [0-0.11] which gathered a high proportion of variants related to the contaminant sample. After removing variants in low-complexity repeats and segmental duplication regions (GnomAD 2.0.2 [Genome Aggregation Database [10]]), the number of germline Single Nucleotide Variants (SNVs) for the contaminated samples was minimized in most of the contamination ratio conditions (including very low contamination ratios).

To determine the detection performance and related threshold in our selected AB range [0-11], we observed the distribution of the number of variants (position only) belonging to the contaminant and the contaminated sample for a range of contamination ratios (between 0% and 50%) (Figure 4). In this distribution, a contamination level of 5% showed an equivalent proportion (55%) of variants belonging to the contaminant and contaminated samples (Figure 4).

**Figure 4.**
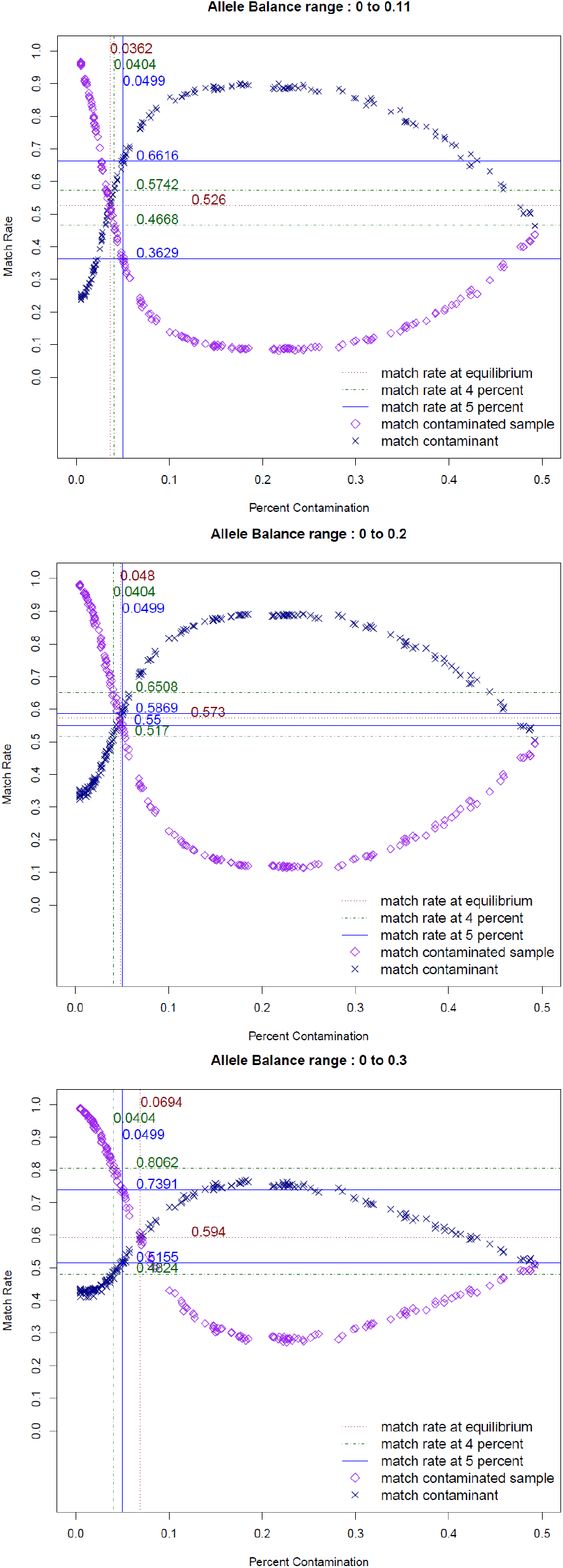
Allele balance range selection for contaminant identification: A) from 0 to 0.11 B) from 0 to 0.2 C) from 0 to 0.3; Blue crosses and purple diamonds represent the evolution of the ratio of variants in the AB range belonging respectively to the initial sample and the contaminant for increasing rate of contamination

A detailed analysis of the curves showed that for contamination ratios under 3.6% the ratio of matching variants related to the contaminant decreased below the proportion of variants related to the contaminated sample. This observation led us to define a minimal threshold of 5% for contaminant identification, to ensure that variants from the contaminant sample were predominantly used for contaminant identification.

Consequently, this contamination threshold of 5% required us to define a shared variant match-rate threshold to validate the contaminant identification. After exploring the impact of contamination by family related samples (Figure 5), we decided to set a minimal 60% match-rate between the selected variant in the AB ranges [0-11%] and the variants in the tested sample. This 60% match-rate threshold supports a clear discrimination between the contaminant and other tested samples as it is 10% above closely related samples.

**Figure 5.**
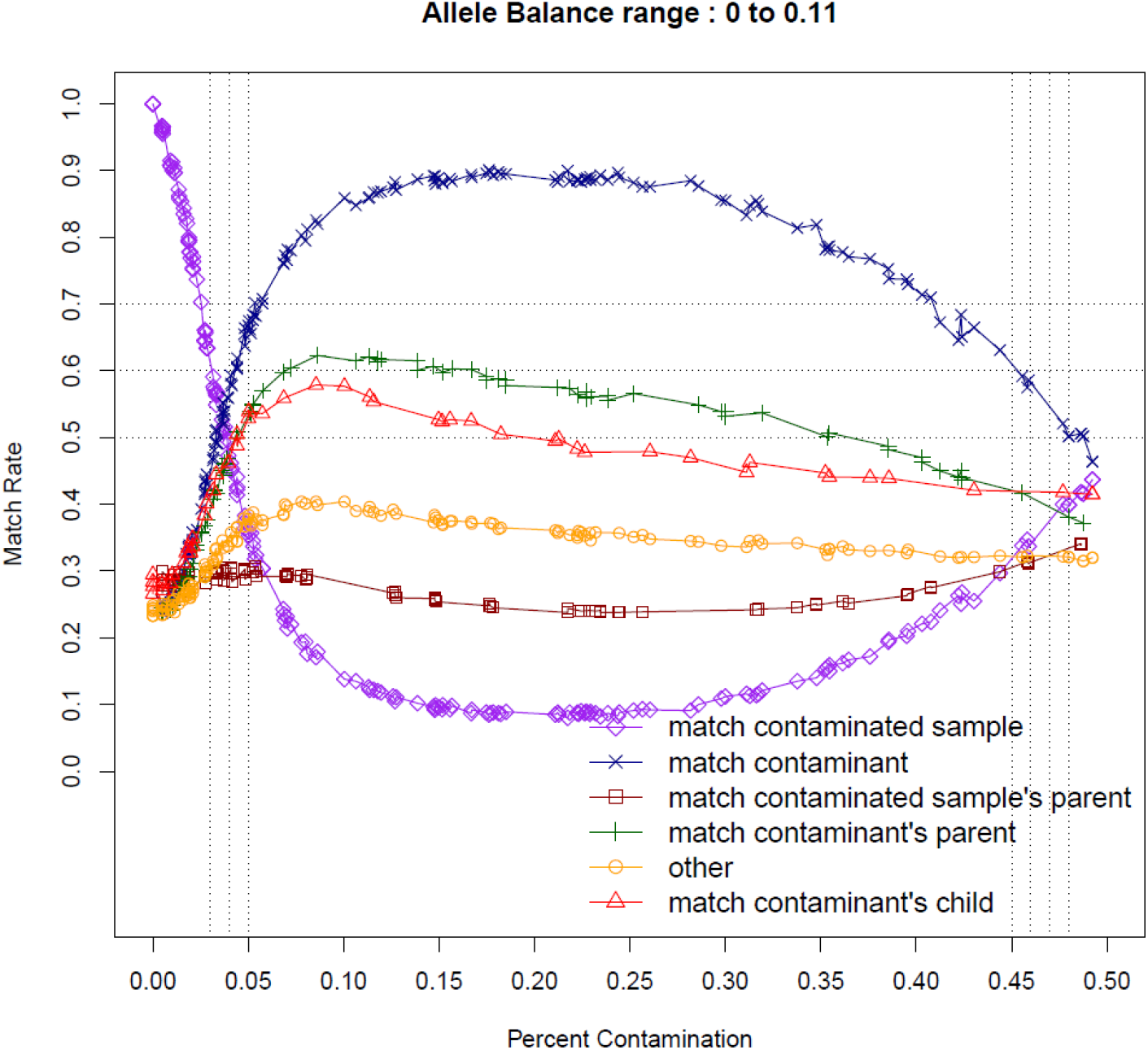
Discrimination between parents and real contaminant in contaminant identification. Blue crosses and purple diamonds represent the evolution of the ratio of variants in the AB range [0-0.11] belonging respectively to the initial sample and the contaminant for increasing rate of contamination. Green crosses and Red triangle represent the same distribution with family related samples. Brown squares and orange round represent cases of unrelated samples

## Results

In order to validate the relevance of our *in silico* simulation, we compared the AB distribution of the experimental mixture against *in silico* mixture conditions. The two distributions shared similar outlines thus validating the guiding principles of this work (Figure 2).

Next, we launched ContaTester and demonstrated contamination level estimations closely approaching those of expected experimental conditions (Table 2). This shows that ContaTester provides correlation and regression as two complementary approaches to estimate the contamination rate. The second order polynomial regression model shows an accuracy down to a mixture prediction of 1%. The results of the Pearson’s correlation are consistent with those of the second order polynomial regression; hence this method can be used for cross-validation and interpretation.

**Table 2.** ContaTester and VerifyBamID2 estimations of contamination ratios from 30x Whole Genome Sequencing of experimental sample mixtures

| Sample mixture contamination level | VerifyBamID2 | ContaTester |  |
| --- | --- | --- | --- |
|  |  | second order polynomial regression | Pearson's Correlation |
| 1.0% | 1.03% | 0.87% | 1.14% |
| 2.5% | 2.50% | 2.19% | 2.75% |
| 5.0% | 4.65% | 4.76% | 4.09% |
| 10.0% | 9.21% | 9.62% | 8.63% |
| 15.0% | 15.49% | 17.17% | 14.71% |

In addition, using BAM files as input, we compared the ContaTester results against VerifyBamID2 and found similar results for contamination estimation in our different experimental mixtures (Table 2). These results show that ContaTester offers a fast and simple alternative for estimating contamination levels, and requires only VCF files.

Lastly, we applied ContaTester’s contaminant detection function to each of the mixture conditions. For contamination levels above 5%, we clearly detected the contaminant NA10859 with more than 60% shared positions within the subset used for the comparison and tested samples (Table 3).

**Table 3.** Contaminant identification with ContaTester from 30x Whole Genome Sequencing sequencing of experimental sample mixtures

| Sample mixture contamination level | Variant count in subset | NA10859 shared variant positions ratio | NA12878 shared variant position ratio |
| --- | --- | --- | --- |
| 1.0% | 13 500 | 0.248 | 0.307 |
| 2.5% | 17 852 | 0.404 | 0.238 |
| 5.0% | 34 918 | **0.691** | 0.127 |
| 10.0% | 89 679 | **0.884** | 0.051 |
| 15.0% | 153 871 | **0.914** | 0.034 |

In summary, ContaTester provides estimations of contamination levels from VCF files generated from germline sample WGS. It provides reliable contaminant identification for cohorts or experimental batches with contamination levels above 5%. ContaTester does not require additional files or parameters, and it provides fast results and reports to support automatic or visual checks of the contamination status. The input, restricted to VCF files (~ 1GB for a 30x WGS), reduces the storage footprint required for quality control compared to solutions that use BAM files (~ 70GB for a 30x WGS). ContaTester requires 70 seconds with a 1 core Intel^®^ Xeon^®^ CPU E5-2680 v4 @ 2.40GHz to process a VCF file obtained from WGS and determine the related contamination level. In comparison, the processing of a BAM file by VerifyBamID2 takes 42 minutes to complete (Table 4). Moreover, contaminant identification by ContaTester takes only 31 minutes for a cohort of 96 samples including 32 contaminated samples (Table 4). In conclusion, ContaTester can be used for the human reference genomes GRCh37 and GRCh38 (Table 2, Table 5) and offers a scalable method for analysis of the increasing volume of VCF files from large WGS projects.

**Table 4.** ContaTester time and memory consumption depending on the number of samples and treatment, on broadwell Intel^®^ Xeon^®^ CPU E5-2680 v4 @ 2.40GHz

| Number of samples | Contaminated samples (> 5%) | Contaminant check | CPU | Parallelization | ContaTester |  | VerifyBamID2 |  |
| --- | --- | --- | --- | --- | --- | --- | --- | --- |
|  |  |  |  |  | Memory (MB) | Time | Memory (MB) | Time |
| 1 | - | - | 1 | 1 | 165 | 1'09" | 536 | 42'11" |
| 96 | - | - | 48 | 48 | 201 | 2'18" | 660 | 1h14'20" |
| 96 | 10 | + | 196 | 48 | 185 | 11'08" | - | - |
| 96 | 32 | + | 196 | 48 | 201 | 30'53" | - | - |
| 900 | - | - | 48 | 48 | 247 | 17'47" | 660 | 9h26'52" |
| 900 | 90 | + | 196 | 48 | 1438 | 12h29'14" | - | - |

**Table 5.** ContaTester results for contamination evaluation of experimental contamination at 30x in Whole Genome Sequencing, aligned on GRCh38

| Sample mixture Contamination level | Correlation | Regression |
| --- | --- | --- |
| 1.0% | 1.14% | 1.64% |
| 2.5% | 2.75% | 2.92% |
| 5.0% | 4.09% | 5.47% |
| 10.0% | 8.63% | 10.25% |
| 15.0% | 14.71% | 17.66% |

## Appendix

### Acronyms

*r*^2^: coefficient of determination. 2
AB: Allele Balance. 2, 3
AD: Allele Depth. 2
BAM: Binary Alignement Map. 1, 3, 4
GnomAD: Genome Aggregation Database. 3
InDels: Insertions-Deletions. 2
SAM: Sequence Alignement Map. 1
SNVs: Single Nucleotide Variants. 3
VCF: Variant Call Format. 1-4
WGS: Whole Genome Sequencing. 2, 4

## Availability of data and materials

Project name: ContaTester

Project home page: https://github.com/CNRGH/contatester

Operating system(s): Runs natively on linux and on any operating system supporting container images (Docker: https://hub.docker.com/r/cnrgh/contatester)

Programming language: Python, R, Bash

Other requirements: Python 3.6 or higher, python libraries (pathlib, os, typing, argparse, io, subprocess, sys, glob, datetime), R 3.3.1, bcftools 1.9 or higher, pegasus 4.8.2 or higher

License: CeCILL

Any restrictions to use by non-academics: None

## Authors’ contributions

D.Delafoy, V.M. conceived the project and drafted the manuscript. D.Delafoy, V.M., F.S., E.L revised the manuscript. D.Delafoy, J.M designed and implemented the code for ContaTester. D.Delafoy, J.M., N.W., T.M. designed and implemented the code for packaging and distribution. D.Delafoy, S.M, D.Daian prepared data and performed test and benchmarking. A.B., R.O. conceived, planned and carried out the experiments. D.Daian, SM carried out sequencing quality control. JF.D. supervised the project as director of the CNRGH. All authors provided critical feedback of the manuscript and read and approved the final manuscript.

## Acknowledgements

We acknowledge the contribution of our colleagues at the CEA - Centre National de Recherche en Genomique Humaine: Elizabeth May and Zuzanna Gerber for careful editing and improvement of the English in this paper, Floriane Lenfant, Marie-Laure Moutet, Feroze Golamaully and Bertrand Fin from “Laboratoire Banque (LB)” for providing experimental mixture samples, Céline Baulard, Florence Jobard, Jeanne-Antide Perrier, David Derbala, Emmanuel Menard, Johann Tassin and Céline Besse from “Laboratoire Plateforme de Production en Génomique Humaine (L2PGH)” for processing and sequencing the samples and Ghislain Septier from the “Laboratoire de Bio-infomatique (LBI)” for ContaTester beta testing. We also acknowledge the CEPH biobank for providing samples.

